## Supplementary figures and images for "Identifying regulators of associative learning using a protein-labelling approach in *C. elegans*"

### Fig S1

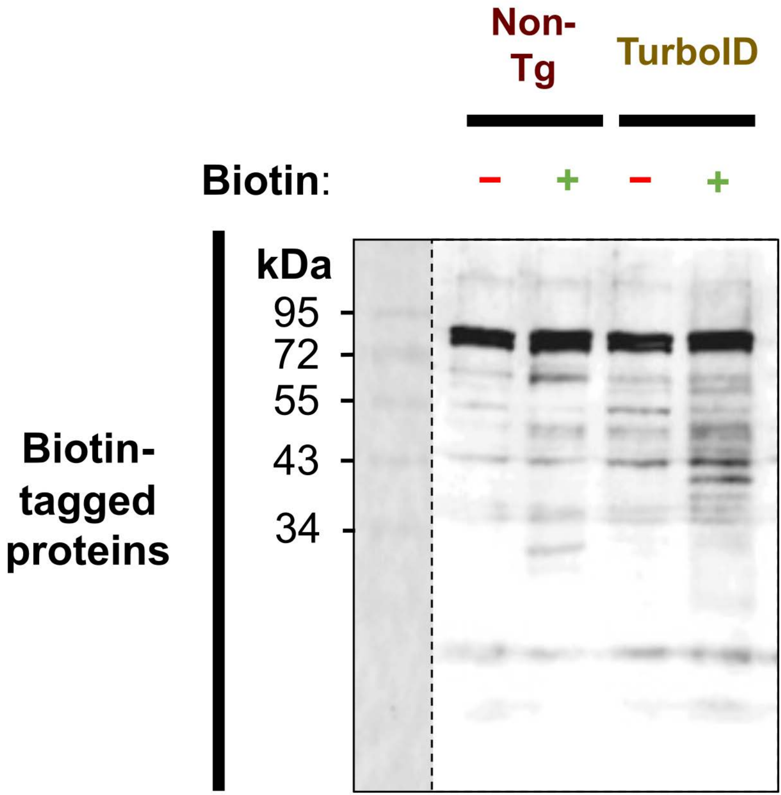

### Fig S2

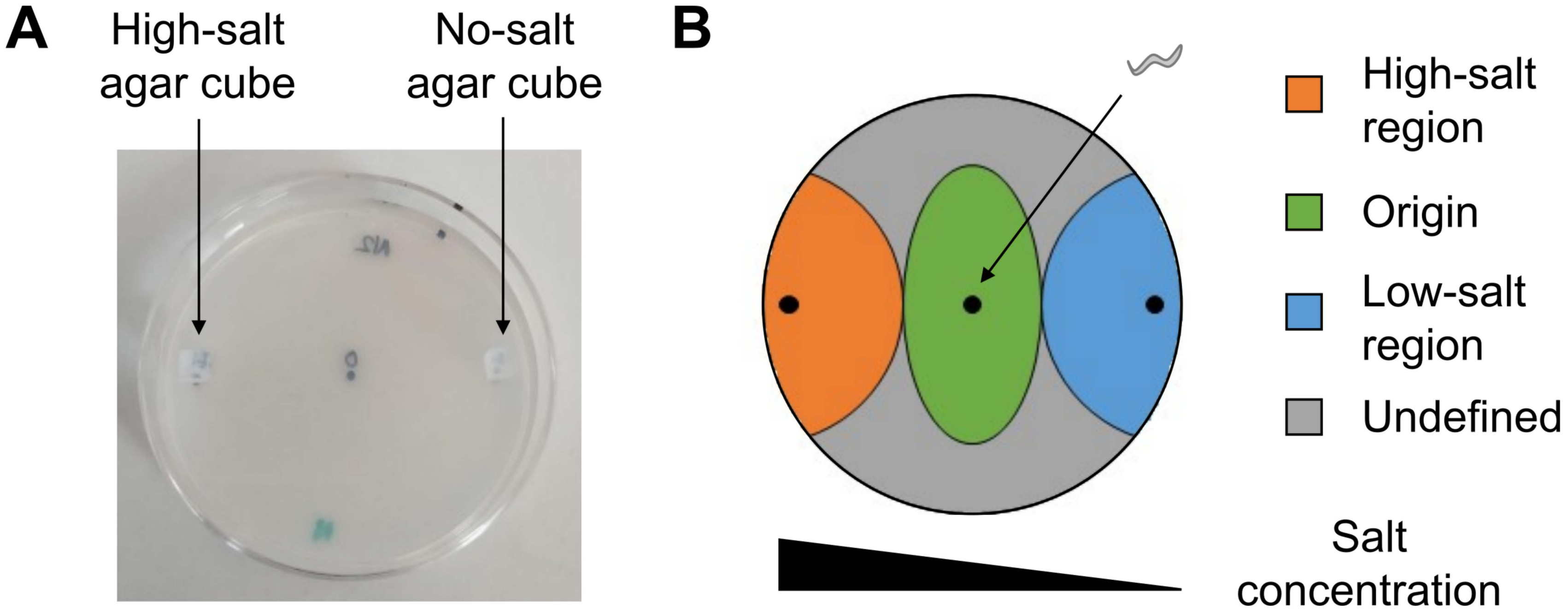

### Fig S3

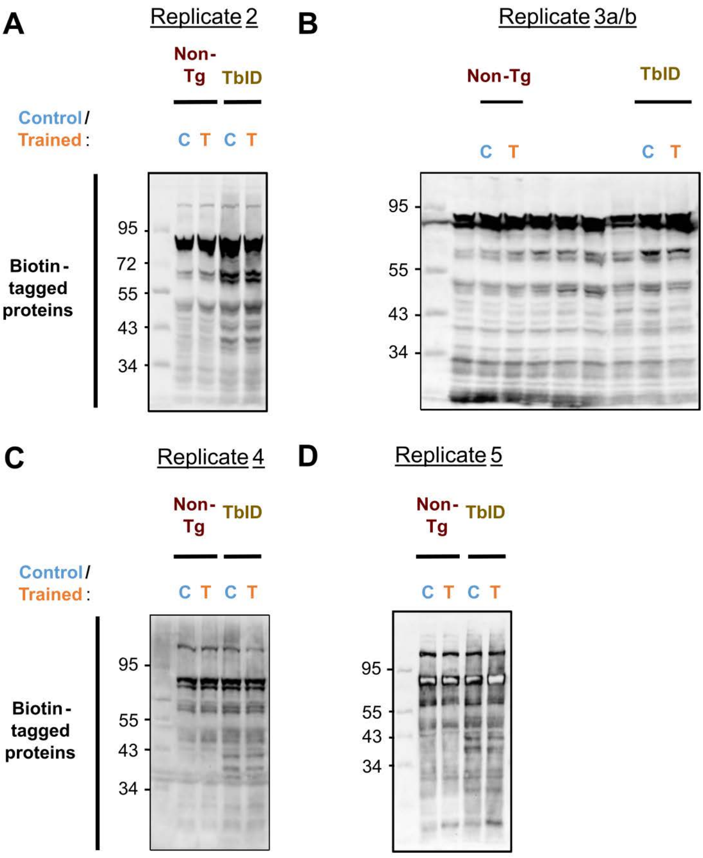

### Fig S4

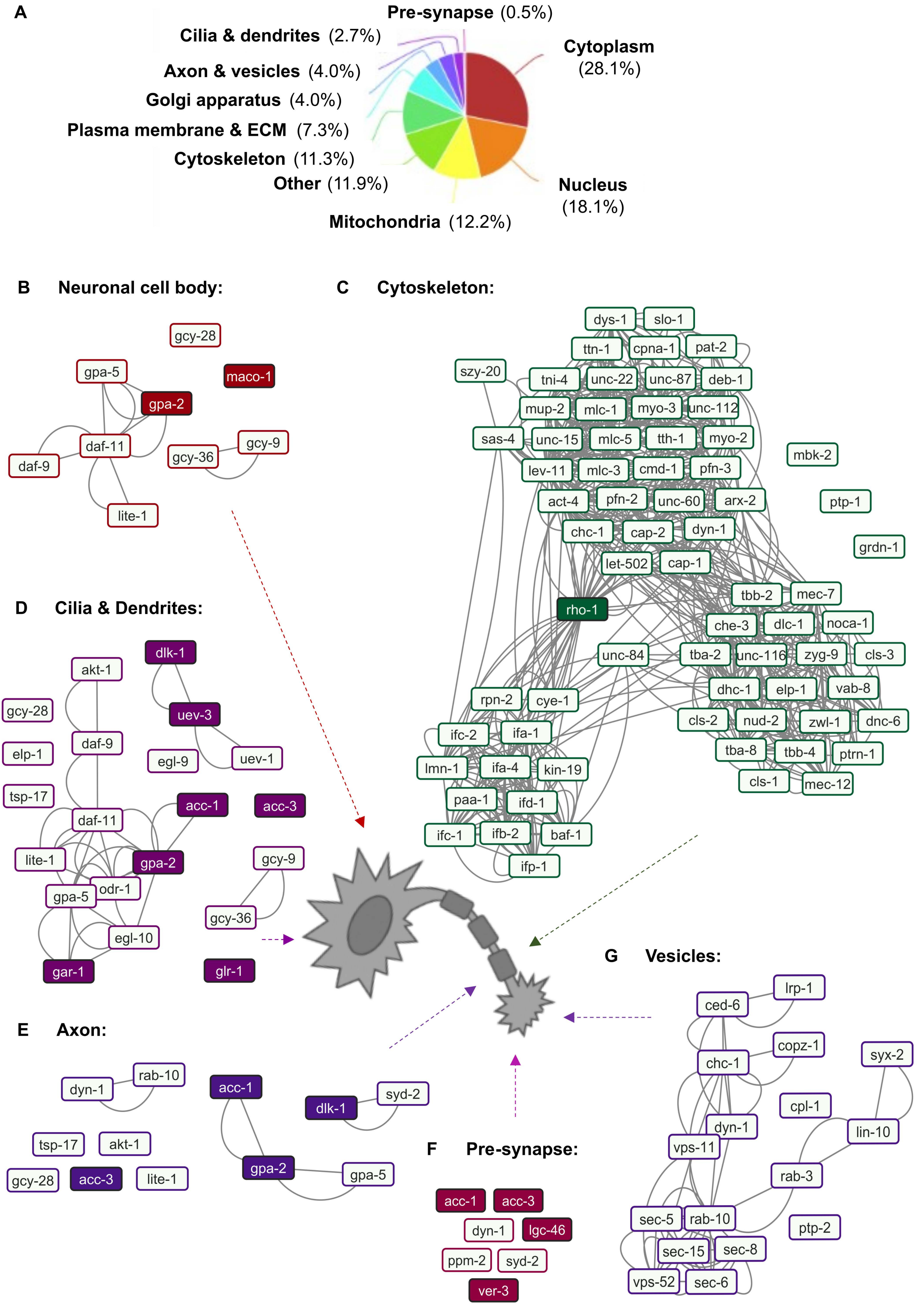

### Fig S5

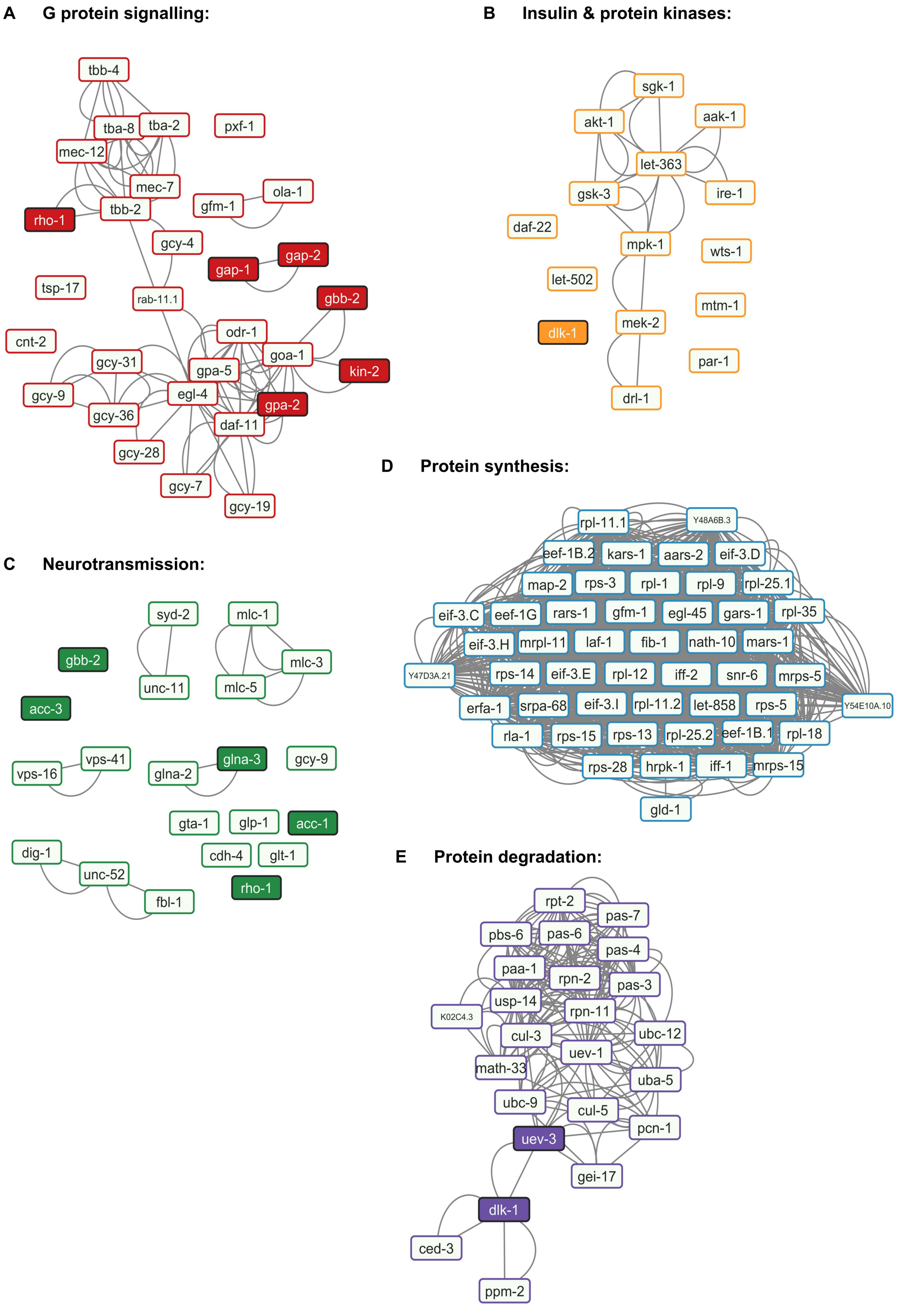

### Fig S6

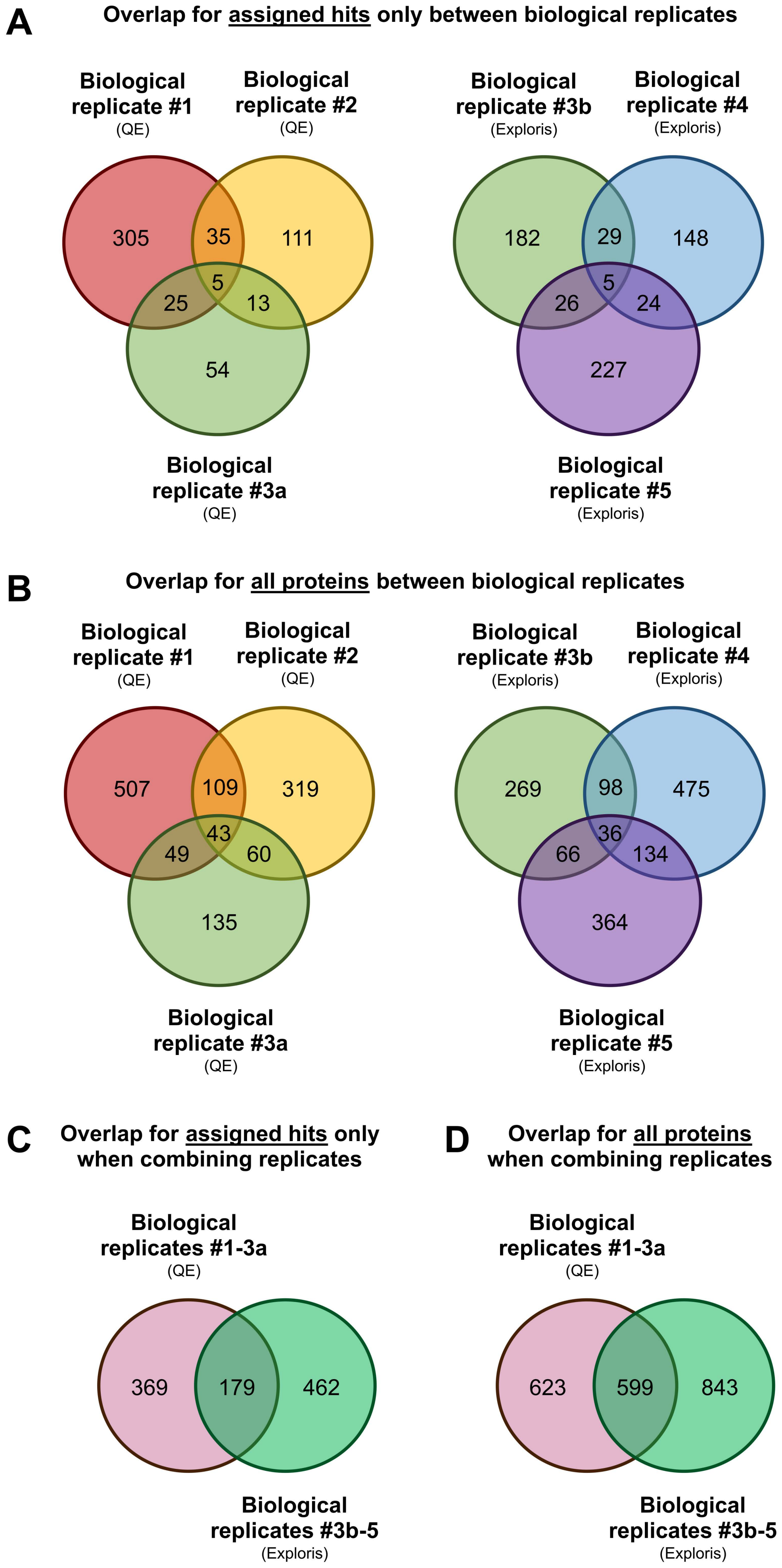

### Fig S7

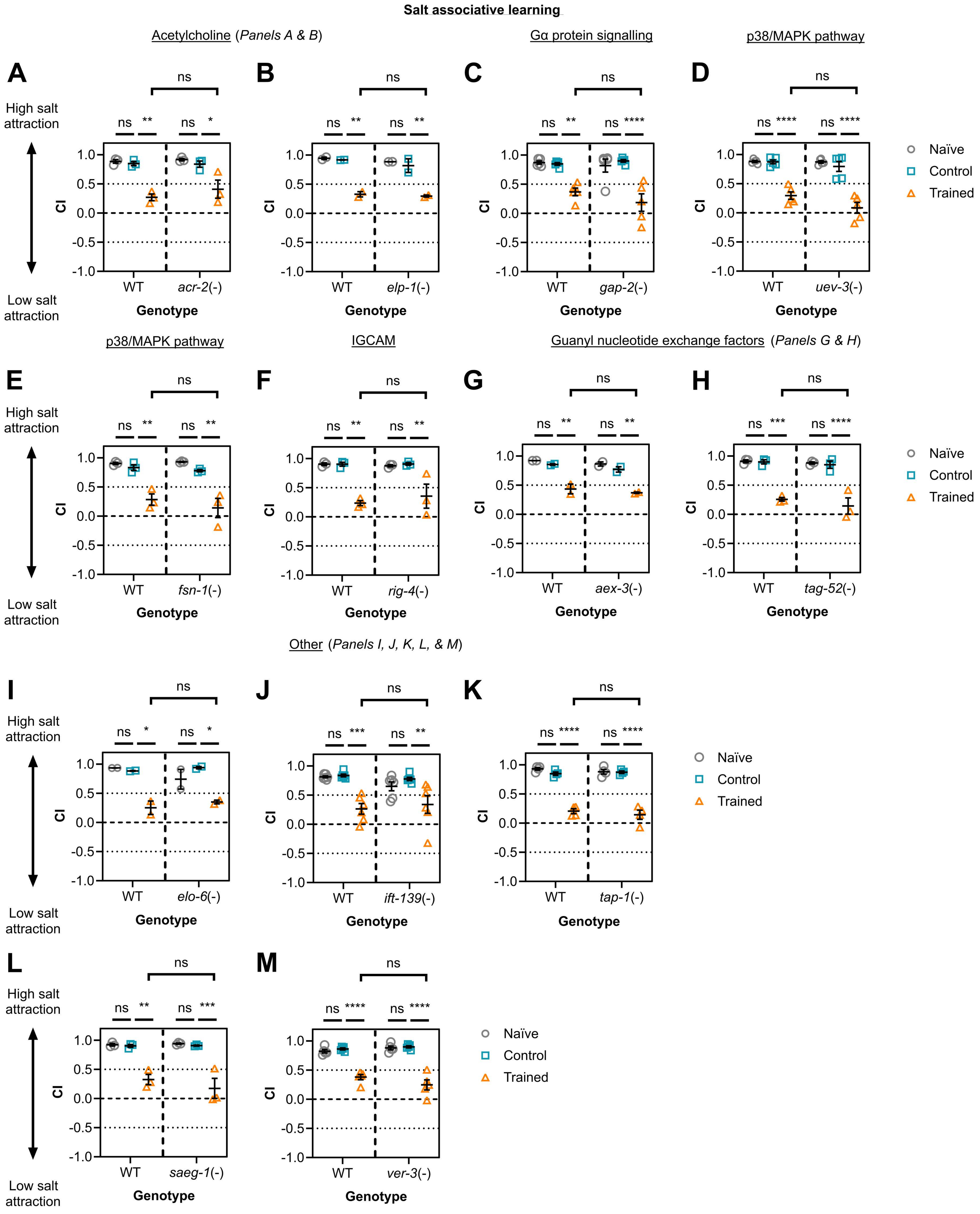

### Fig S8

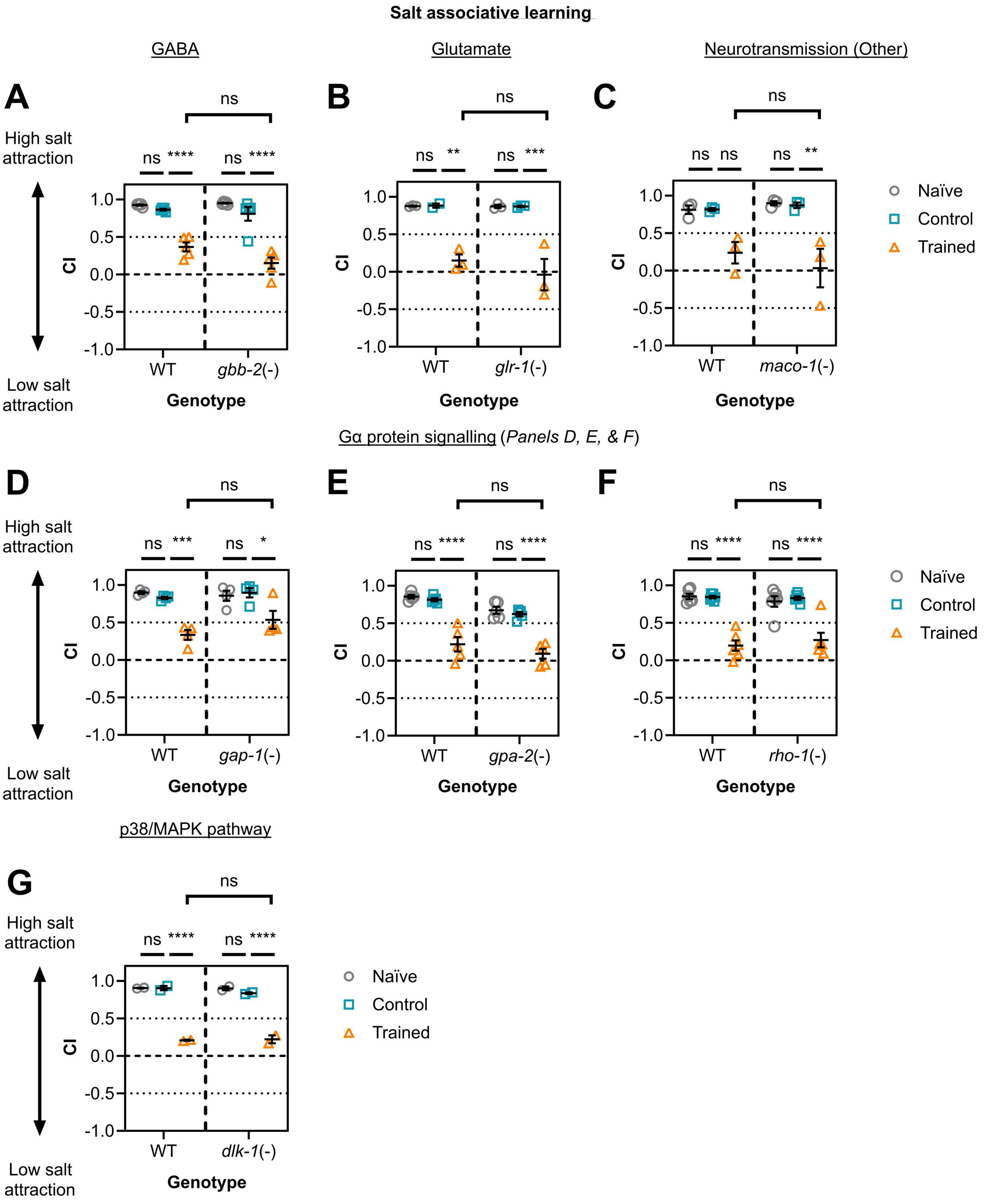
